# Deep graph learning of inter-protein contacts

**DOI:** 10.1101/2021.08.14.456342

**Authors:** Ziwei Xie, Jinbo Xu

**Affiliations:** Toyota Technological Institute at Chicago, Chicago, IL 60637, USA

## Abstract

**Motivation:** Inter-protein (interfacial) contact prediction is very useful for *in silico* structural characterization of protein-protein interactions. Although deep learning has been applied to this problem, its accuracy is not as good as intra-protein contact prediction.

**Results:** We propose a new deep learning method GLINTER (Graph Learning of INTER-protein contacts) for interfacial contact prediction of dimers, leveraging a rotational invariant representation of protein tertiary structures and a pretrained language model of multiple sequence alignments (MSAs). Tested on the 13th and 14th CASP-CAPRI datasets, the average top L/10 precision achieved by GLINTER is 54.35% on the homodimers and 51.56% on all the dimers, much higher than 30.43% obtained by the latest deep learning method DeepHomo on the homodimers and 14.69% obtained by BIPSPI on all the dimers. Our experiments show that GLINTER-predicted contacts help improve selection of docking decoys.

## 1 Introduction

Proteins perform functions by interacting with other molecules or forming protein complexes. As a result, the full characterization of protein-protein interactions with structural details is crucial to atom-level understanding of protein functions. The *in silico* structural characterization of protein complexes, or quaternary protein structure prediction, is a long standing challenge in computational structural biology. Given individual protein chains (and possibly their structures), interfacial contact prediction aims to predict which pairs of residues on the protein surface are geometrically close to each other after the protein chains bind together. Interfacial contacts may facilitate generating and filtering docking decoys^1,2,3,4^, and reveal important biophysical properties and evolutionary information of protein interfaces^5^. They are also useful for the redesign of protein-protein interfaces^6^ and prediction of binding affinity^7^.

Co-evolution analysis by global statistical methods^8,9^ has been used for inter-protein contact prediction. A recent study^10^ showed that co-evolution-based *in silico* protein-protein interaction screening methods produced more true protein-protein interactions than high-throughput experimental techniques. Nevertheless, accurate co-evolution analysis needs a large number of sequence homologs and thus, may not work well on a large portion of heterodimers for which it is very challenging to find sufficient number of interacting paralogs (interlogs)^11–13^. On the other hand, protein language models, which are trained on individual protein sequences or MSAs (multiple sequence alignment), are shown to perform similarly as or better than global statistical methods on intra-chain contact prediction when few sequence homologs are available^14,15^. We hypothesize that the language models trained on individual protein chains may also generalize to protein-protein interactions, reducing the required number of interlogs. Protein language models are also much faster since they require only one-time forward computation during inference and thus, more suitable for proteome-scale screening of protein-protein interactions.

RaptorX-ComplexContact^11,16^ possibly is the first supervised deep learning method for interfacial contact prediction. It is mainly developed for heterodimers, although can be used for homodimers. Nevertheless, its deep models are purely trained on individual protein chains instead of protein complexes. Further, RaptorX-ComplexContact does not make use of any (experimental or predicted) structures of constituent monomers of a dimer. Recently some deep learning methods are developed specifically for contact prediction of a homodimer, e.g., DNCON_inter^17^ and DeepHomo^18^, both using ResNet originally implemented in RaptorX^19^. In addition to evolution information, DeepHomo uses docking maps, native intra-chain contacts, and experimental structural features derived from monomers to achieve state-of-the-art performance. However, it is slow in calculating docking maps and thus, cannot scale well to proteome-scale prediction. Some deep learning methods also use learned representations of tertiary structures, including voxels^20,21^ and radial/point cloud representations on protein surfaces^22–24^. Meanwhile, some representations include anisotropy information in the structures^25,26^ while others do not.

Given the tremendous progress in protein structure prediction^19,27–30^ and the fast growing number of protein sequences, it is important to leverage predicted structures of constituent monomers and large sequence corpus to produce accurate, proteome-scale interfacial contact predictions. An interfacial contact prediction method shall effectively extract coevolution signals from a small number of interlogs, and make use of predicted structures of constituent monomers. Here we propose a new supervised deep learning method GLINTER for interfacial contact prediction that integrates representations learned from (experimental and predicted) monomer structures and attentions generated by the MSA Transformer (ESM-MSA)^14^ from interlogs of the dimer under prediction. GLINTER applies to both heterodimers and homodimers, outperforming RaptorX-ComplexContact, DeepHomo and BIPSPI on the 13th and 14th CASP-CAPRI datasets. The contacts predicted by GLINTER may also improve the ranking of the HDOCK-generated docking decoys^31^. Further, our method runs very quickly, which makes it suitable for proteome-scale study.

## 2 Methods

### 2.1 Network Architecture

As shown in Fig. 1, our network, denoted as GLINTER, consists of two major modules: a Siamese graph convolutional network (GCN)^32^ and a 16-block ResNet^33^. The GCN extracts local features from 3 types of graphs derived from monomer structures. The ResNet takes as input the outputs of the GCN module and the attention weights generated by the MSA Transformer^14^ and yields interfacial contact prediction. One ResNet block has two convolutional layers, each with 96 filters and a 3×3 kernel. ELU and BatchNorm are used in each block. ResNet is connected to a fully connected layer and a softmax layer for contact probability prediction.

**Fig. 1.**
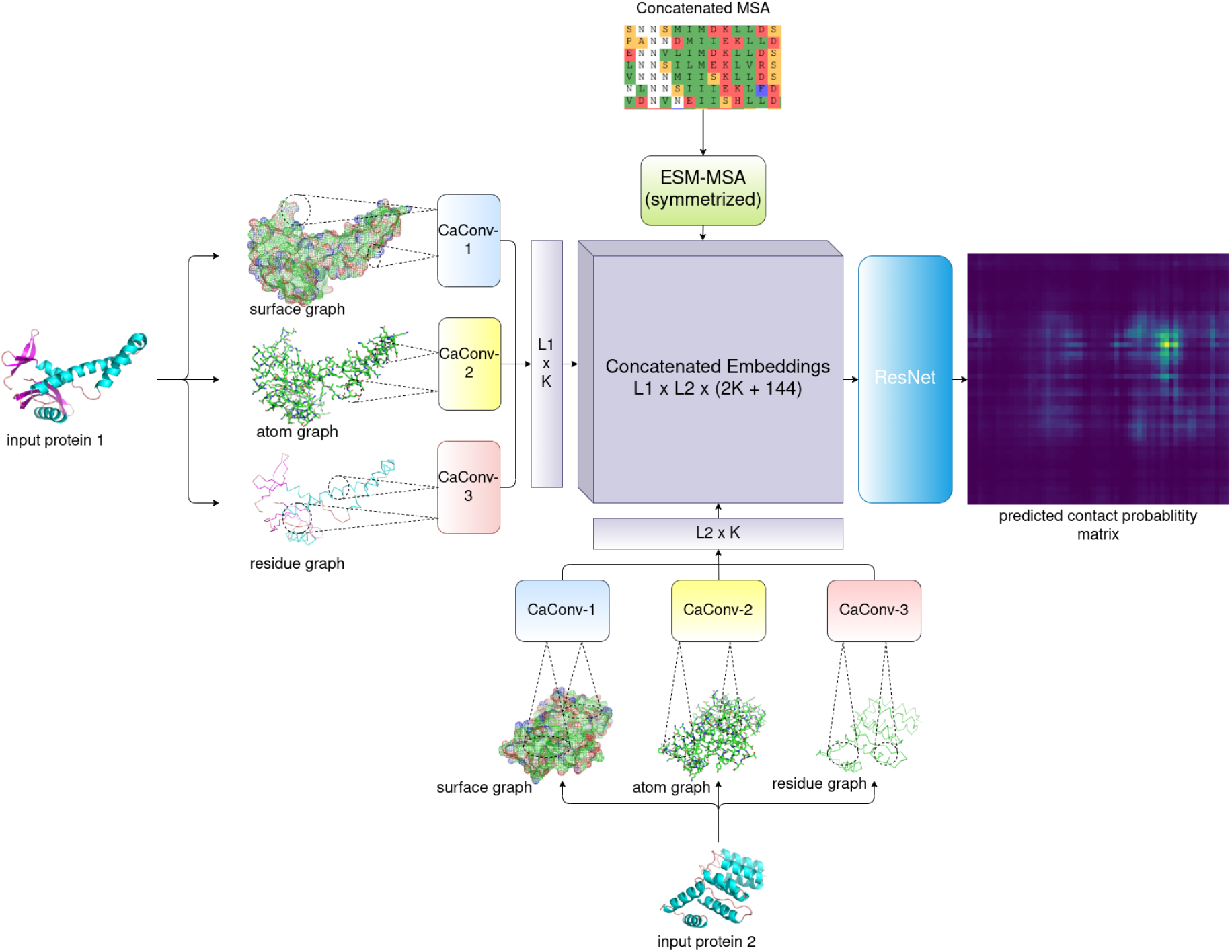
Overview of the GLINTER architecture. L1 and L2 are the lengths of the two protein chains, K is the number of channels in a CaConv layer, and 144 is the total number of heads in the row attention weights generated by Facebook's MSA Transformer.

At each graph convolution layer (denoted as CaConv), we calculate the message for a graph edge and node as follows. For an edge, we feed its feature and the features of its two ends to a subnetwork to generate a message. For a node *q*, we first aggregate all messages of its adjacent nodes using max pooling, and then pass the result to a subnetwork to generate a message of *q*, i.e.

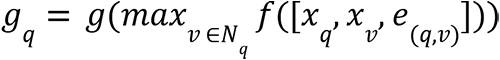

where *x_q_* is the feature of node *q*, *ν* is a node in the neighborhood *N_q_* of*q*,*x_ν_* is the feature of *ν*, *e*_(*q*,*ν*)_ is the feature of edge (*q*, *ν*) and the nonlinear functions *g* and *f* are two fully connected layers of 128 hidden units with BatchNorm and ReLU.

Both coordinates and normals are used to represent the geometric properties of a monomer structure^23^. We standardize the geometric features so that they are invariant to the coordinate system used by the monomer structure. While calculating an message for any node *q* (i.e., computation of *f*), all the adjacent nodes of *q* are first translated using *q* as the origin, and then rotated using its predefined local reference frame^34,35^. The standardized features are then concatenated with other features to form the actual inputs of function *f*.

We use a separate graph convolution network (GCN) module to process each graph. When multiple graphs are used for a monomer, the outputs of all its GCN modules are concatenated to form a single output vector of this monomer. The outputs of two monomers are then outer-concatenated to form a pairwise representation of this dimer. When the ESM row attention weight is used, the attention matrix generated by Facebook's MSA Transformer is concatenated to the pairwise representation, which is then fed to the ResNet for interfacial contact probability prediction.

### 2.2 Features

#### Graph representation of protein structures

We build three different graphs from one protein structure: residue graph, atom graph and surface graph. In a residue graph, a node is a residue represented by its CA atom, and there is an edge between two residue nodes if and only if the Euclidean distance between their CA atoms is within a certain cutoff, e.g. 8Å. In an atom graph, a node is a heavy atom or a residue represented by its CA atom, and there is one edge between one residue node and one atom node if and only if their Euclidean distance is within a certain cutoff.

We use Reduce^36^, MSMS^37^ and trimesh^38^ to construct the triangulated surface of a protein structure (details are in the supplemental file). The surface can be essentially interpreted as a mesh enclosing the protein. Two neighboring triangles in the surface share either one edge or at least one vertex. In a surface graph, one node represents one residue or one vertex on the triangulated surface. We add one edge between one residue node and one triangle vertex if and only if their Euclidean distance is within a certain cutoff. It takes only a few seconds to build a surface graph and thus, our method scales well on large-scale prediction^10^. In contrast, DeepHomo uses a computationally-intensive docking program to generate docking maps.

#### Features

Table S2 summarizes all the features. The geometric features of a residue node include its coordinates and a local reference frame derived from the N-CA-C plane. As shown in Fig. S3, it uses the CA-C bond as the x-axis, the vector perpendicular to the plane formed by the N-CA and CA-C bonds as the z-axis, and their cross-product as the y-axis. Such a representation is rotation invariant and thus, may generalize well without unnecessary data augmentation. The other features of a residue node include PSSM (position-specific scoring matrix), residue solvent accessible surface areas (which summing up the solvent accessible surface areas of all atoms in the residue), the one-hot encoding of amino acid type, and the sequence index of the residue divided by the protein sequence length (which is used to provide order information for neural network architectures that are order invariant)^29^.

In an atom graph, an edge has a binary feature called “edge type”. It is equal to 1 if the nodes of this edge belong to the same residue. An atom is encoded by a 10-dimensional one-hot vector, indicating 4 backbone atom types (CA, N, C, O) and 6 side chain atom types (CB, C, N, O, S, H).

In a surface graph, we use the coordinates and normals generated by MSMS as the features of a triangle vertex^22^, which indicate the contour and orientation of some local patches on the surfaces. Normals are initially computed by MSMS, and then reweighted by trimesh.

#### Coevolution signals generated by Facebook's MSA Transformer

We use the row attention weights generated by the MSA Transformer as interfacial co-evolution signals. We build a joint MSA for a heterodimer using the protocol proposed by RaptorX-ComplexContact^11^. For a homodimer, we simply concatenate each sequence in the MSA with itself. We then select a diverse set of sequences from the joint MSA as the input of the MSA Transformer (detailed in the supplemental file). We further symmetrized the generated inter-chain attentions^14^.

### 2.3 Datasets

#### CASP-CAPRI data

We use all 32 dimers (23 homodimers and 9 heterodimers) with at most 1000 residues in the 13th and 14th CASP-CAPRI datasets^41^ as our test set. We do not include the dimers with more than 1000 residues since Facebook's MSA Transformer cannot handle such a large protein. To avoid redundancy between our training and test sets and to fairly compare GLINTER with recently published methods, we do not use the 11th and 12th CASP-CAPRI data. We run HHblits on the “uniclust30_2016_09” database to build MSAs for individual chains and then concatenate two MSAs to form a joint MSA for a heterodimer using the method described in^11^. We use monomer (bound) experimental structures as inputs since their unbound structures are unavailable. We also tested the 3D structure models of individual chains predicted by AlphaFold^42,43^ in CASP13 and 14, except for T0974s2 which did not have a predicted 3D model.

#### 3 DComplex data

Our training set has 5306 homodimers and 1036 heterodimers derived from 3DCompex^44^. We do not include the dimers with more than 1000 residues due to MSA Transformer's restriction. We say two dimers are at most *x*%similar, if the maximum sequence identity between their constituent monomers is no more than *x*%. A dimer is treated as a homodimer if at least 90% of its two constituent monomer sequences can be aligned with 90% sequence identity; otherwise a heterodimer. We build the training and validation set as follows.

1. Exclude the dimers in 3DComplex similar to the test set, judged by MMseqs2 E-value < 1.
2. Cluster all dimers using the 40% sequence identity threshold. In each cluster, only keep the dimer with the largest number of contacts to reduce redundancy.
3. Randomly choose 100 dimers (82 homodimers and 18 heterodimers) to form the validation set and use the remaining 6342 dimers to form the training set.
4. Finally, run HHblits on the “uniclust30_2016_09” database to generate MSAs for single chains and build a joint MSA as described in the previous subsection.

### 2.4 Training and evaluation

We use weighted cross-entropy as the loss function since the percentage of contacts is very small. The weight of a contact is 5 times that of a non-contact. We trained our deep models using Adam as the optimizer^45^, with the hyperparameters β_1_ = 0. 9, β_2_ = 0. 999, ϵ = 1*e*^−8^;. The learning rate is initialized to 0.0001 and reduced by half every 4 epochs. All models are trained for 20 epochs on 2 Titan X GPUs, with minibatch size 1 on each GPU. It takes 20-40 minutes to train one epoch. For a given hyperparameter setting, we select the model with the best top-10 precision on the validation data as the final model.

Since our deep network is rotation invariant, we do not augment the training set by rotating a monomer multiple times. Nevertheless, we randomly rotate a monomer once before training to prevent our deep network from learning unexpected artifacts in the data set. For a heterodimer, we use both of the orders of its two proteins in training. For evaluation, we predict two contact maps for one heterodimer by exchanging the order of its two proteins, and then compute the geometric average of the two predicted contact map probability matrices as the final prediction.

We evaluate contact prediction in terms of AUC (area under curve) and top k precision where k=10, 25, 50, L/10 and L/5 and L is the length of the shorter protein in a dimer. When the number of native contacts is less than k, we still use k as the denominator while computing the top k precision. Inter-chain contact maps are more sparse than intra-chain contact maps, so we evaluate a smaller number of predicted inter-chain contacts.

### 2.5 Methods to compare

We compare GLINTER with DeepHomo, RaptorX-ComplexContact and BIPSPI. DeepHomo is a ResNet-based method developed for only homodimers. RaptorX-ComplexContact is a sequence-only and ResNet-based method developed mainly for heterodimers. Both DeepHomo and RaptorX-ComplexContact take as input the coevolution information computed by CCMpred^39^ while GLINTER does not. BIPSPI works for both homodimers and heterodimers and can take both structures and MSAs as input.

## 3 Results

We test our method with the bound experimental structures while comparing it with BIPSPI and DeepHomo. We also study the impact of the quality of predicted structures on our method. Following DeepHomo^18^, we say there is one true contact between two residues (of two monomers) if in the experimental complex structure the minimal distance between their respective heavy atoms is less than 8Å.

### 3.1 Evaluation of interfacial contact prediction

As shown in Table 1, on the 23 test homodimers, GLINTER has 54.35% top 10 precision and 51.16% top L/10 precision, while DeepHomo has 30.43% top 10 precision and 27.32% top L/10 precision. Tested on the 9 heterodimers, GLINTER has 44.44% top 10 precision and 48.32% top L/10 precision, while ComplexContact has 14.44 % top 10 precision and 13.58% top L/10 precision. Even using the monomer structures predicted by AlphaFold-1 and AlphaFold-2 as input, GLINTER has 43.04% top 10 precision on the homodimers and 24.44% top 10 precision on the heterodimers. In summary, GLINTER consistently outperforms DeepHomo and ComplexContact by a large margin no matter whether experimental or predicted monomer structures are used.

**Table 1.**
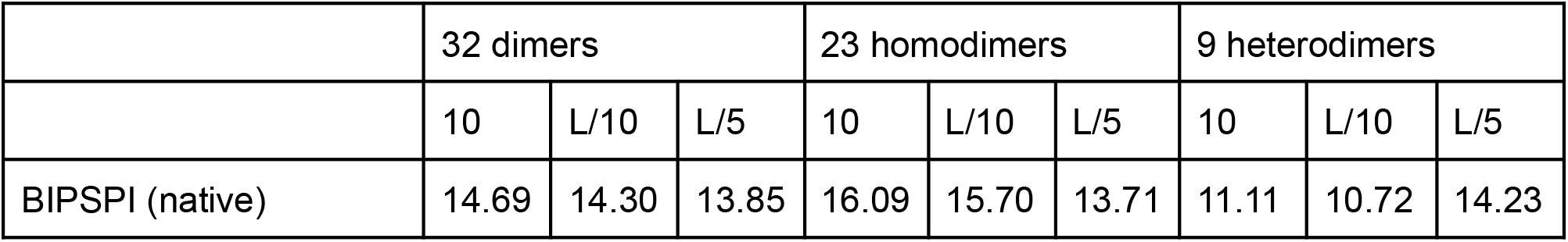
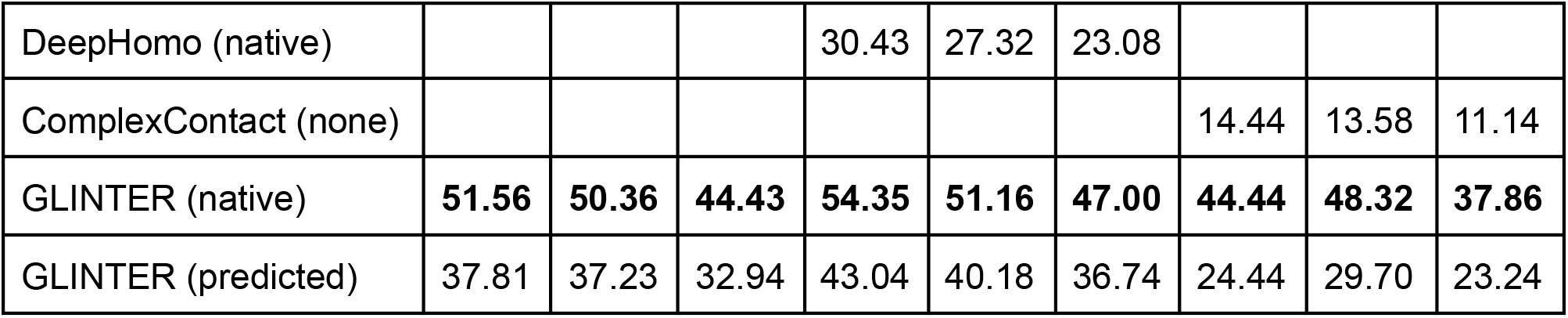
Average contact prediction precision on the CASP-CAPRI test sets. “Native” means the experimental monomer structures are used. “None” means that tertiary structures are not used at all. “Predicted” means the monomer structures predicted by AlphaFold are used.

### 3.2 Ablation study

We train the GLINTER models under 8 different settings (different sets of input features). Tables S3 and S4 show their test results with monomer experimental structures and AlphaFold-predicted monomer structures, respectively. We have studied the following 8 settings: “Residue”, “Residue+ESM”, “Residue+Atom”, “Residue+Atom+ESM”, “Residue+Surface”, “Residue+Surface+ESM”, “Residue+Atom+Surface'’ and “Residue+Atom+Surface+ESM” models. Here “Residue”, “Atom” and “Surface” represent the residue, atom, and surface graphs, respectively. “ESM” means that the ESM row attention weights are used. Using the ESM row attention weights does not change the network architecture, but increases the input dimension of the first ResNet block, as shown in Fig 1.

To evaluate the contribution of the ESM row attention weights, we test a baseline model called ESM-Attention that uses only the ESM row attention weights as input. As shown in Fig. S1, the major module of this model is a 2D ResNet with the same architecture as the one used in the Residue+ESM model.

To evaluate the contribution of the graph convolution module, we develop a baseline model denoted as “CNN+ESM-Attention”, which uses an 1D convolutional network (CNN) and the same set of input features. Similar to the Residue+ESM model, the CNN+ESM-Attention model consists of two major modules: a Siamese 1D CNN and a ResNet. The 1D CNN has 4 convolution layers (each with 128 filters and kernel size 5) and the ResNet is the same as that used in the Residue+ESM model (Fig. S2). Both the ESM-Attention and the CNN+ESM-Attention models are trained on the same dataset using the same protocols as the GLINTER models.

#### Contribution of the graph convolution module

As shown in Table S3, the CNN+ESM-Attention model has similar performance as the ESM-Attention model. The best CNN+ESM-Attention model has 35.31 % top-10 precision and 24.31% top-L/10 precision, while the ESM-Attention model has 30.93% top-10 precision and 29.38% top-L/10 precision. In contrast, the Residue+ESM model has 42.81% top-10 precision and 41.80% top-L/10 precision, which suggests that the residue graph (derived from monomer structures) used by GLINTER is indeed very helpful for interfacial contact prediction.

#### Dependency on distance cutoff

The distance cutoff used to define graph edges is an important hyperparameter. According to our observation, a model with a larger distance cutoff tends to have a lower training loss, although its prediction performance may not be as good. A model with a smaller distance cutoff may have a higher training loss and much worse prediction performance. As shown in Table S3, the top k precision of GLINTER models increases along with the distance cutoff until reaching the optimal value. For example, the top-10 precision of the Residue+Atom model increases from 21.56% to 32.50% as the distance cutoff increases from 4Å to 6Å, and then decreases to 26.56% when the distance cutoff is 8Å. This saturation effect on the distance cutoffs is also observed in^20^.

Different types of graphs may rely on distance cutoffs differently. For example, the top 10 precision of the Residue+Surface model is around 33% when the distance cutoff defining the surface graph ranges from 4Å to 10Å, while the precision of the “Residue+Atom” model changes a lot with respect to the distance cutoff. Here we determine the optimal distance cutoff using the experimental monomer structures, which may not have the optimal performance when predicted monomer structures are used.

#### Dependency on the quality of predicted monomer structures

GLINTER models are trained with monomer experimental structures. Here we study their prediction performance when the AlphaFold-predicted monomer structures are used. We use the lower TMscore^46^ of the two constituent monomer models to measure the structure quality of a dimer under test. We exclude the test dimers without any correct top k predicted contacts when their native structures are used as input. Since there are only dozens of test targets, we divide them into four groups according to their TMscores: low quality (0.2≤TMscore<0.5), acceptable quality (0.5≤TMscore<0.7), medium quality (0.7≤TMscore<0.9) and high quality (0.9 ≤TMscore<1.0).

Fig. S6 shows that even trained on bound experimental structures, our methods work well on predicted structures with medium or high quality (i.e., TMscore>0.7). When the predicted monomer structures have lower quality (TMscore<0.7), GLINTER models perform better with experimental structures than predicted structures. By comparing Fig. S6(D) and S6(E), we find that the ESM row attention weight may not be able to reduce the precision gap incurred by predicted structures. This suggests that the ESM row attention weight derived purely from MSAs may not necessarily improve the robustness of our structure-based models.

#### Contribution of the ESM row attention weight

As shown in Tables 1 and S3, on the 32 dimer targets, the ESM-Attention model has top 10 and L/10 precision 30.93% and 29.38%, respectively, greatly outperforming BIPSPI, which has top 10 and L/10 precision 14.69% and 14.30%, respectively. That is, even though the MSA Transformer is pre-trained with the MSAs of single-chain protein sequences, it works for inter-chain contact prediction. Over the 9 heterodimer targets, the top 10 precision of ComplexContact and ESM-Attention is 14.44% and 27.78%, respectively. As shown in Tables S3 and S4, no matter whether native or predicted monomer structures are used the ESM row attention weight consistently improves the performance of GLINTER models, which confirms that coevolution signals are very useful for inter-chain contact predictions.

Fig. 2(A) compares the performance of the ESM-Attention model (which is a sequence-only model) and the Residue+Atom+Surface model (which is a structure-only model) when the native structures are used. They have similar overall performance, but perform very differently on individual test targets, which suggests that the ESM row attention weight and structure information are highly complementary to each other. On the majority of test targets, the Residue+Atom+Surface+ESM model outperforms the ESM-Attention model (Fig. 2(B)) and the Residue+Atom+Surface model (Fig. 2(C)). Detailed performance comparison among the three models can be found in Table S1. A case study on target T0997 can also be found in the supplemental file.

**Fig. 2.**
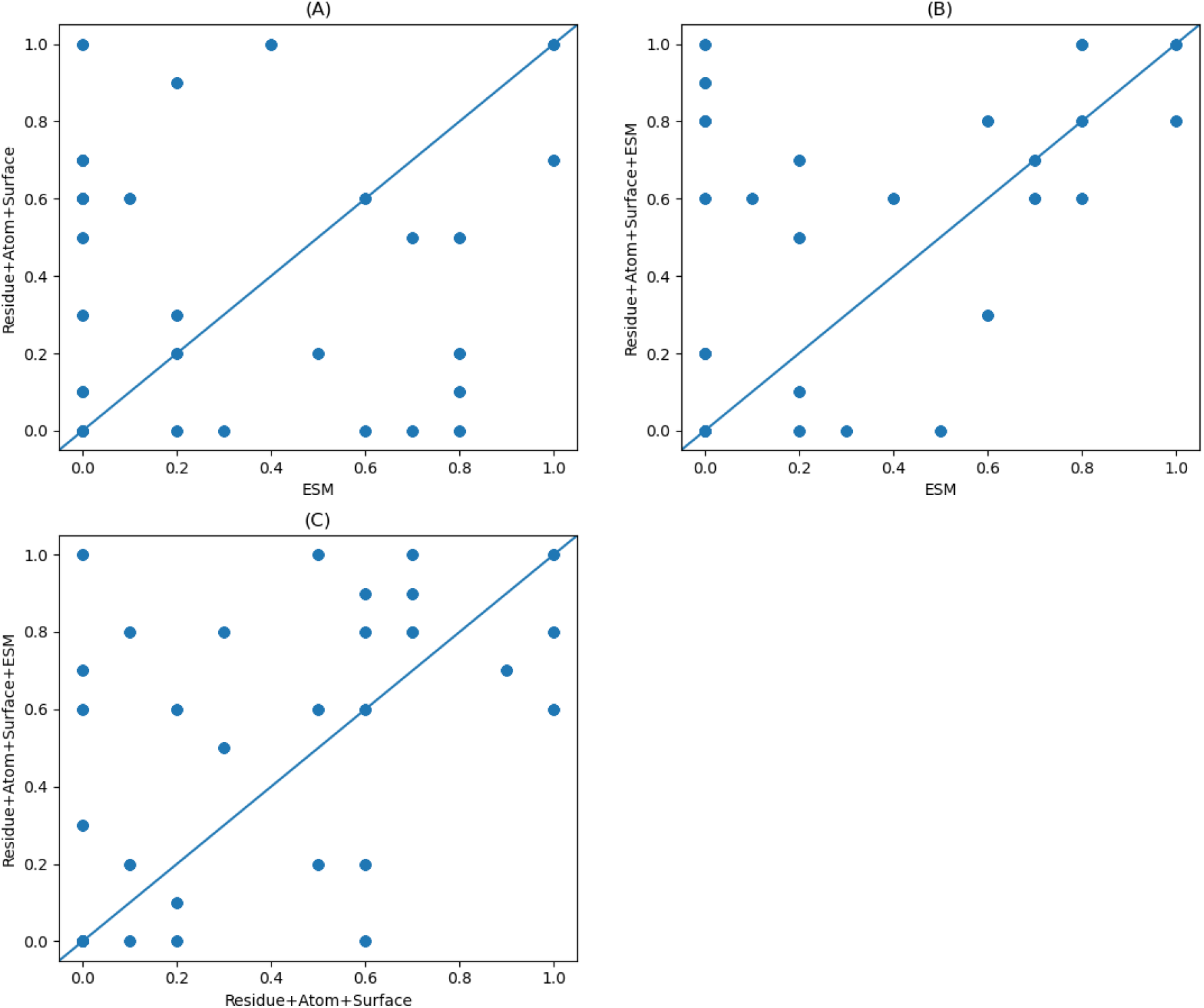
Comparison of top-10 precision of different models.

#### Dependency on the depth of multiple sequence alignments (MSA)

It is known that intra-chain contact prediction precision correlates with the depth of MSAs denoted as Meff (defined in Supplemental File). Here we study the impact of MSA depth on interfacial contact prediction when the ESM row attention weight is used. To remove the impact of inaccurate predicted structures, here we test GLINTER models with native monomer structures. Fig. S7 shows that there is certain correlation (R^2^=0.3093) between the number of correct top-10 predictions by the ESM-Attention model and the ln(Meff) of the input MSA.

### 3.3 Application to selection of docking decoys

A simple application of predicted interfacial contacts is to select the docking decoys. We use the top k (k=10, 25, 50) contacts predicted by the Residue+Atom+Surface+ESM model to rank the docking decoys generated by HDOCK. The quality of a docking decoy is calculated by comparing it with its experimental complex structure using MMalign^47^. For each target, we select top N decoys ranked by the predicted interfacial contacts and define their highest TMscore as the “TMscore of the top N decoys”. In Fig. 3, the y-axis shows the average TMscore of the top N decoys of all the test dimers. Generally speaking, predicted contacts may improve the quality of top decoys by 5-8%. Except when N≤10, generally speaking using more top predicted contacts may select better decoys than using only top 10 predicted contacts.

**Fig. 3.**
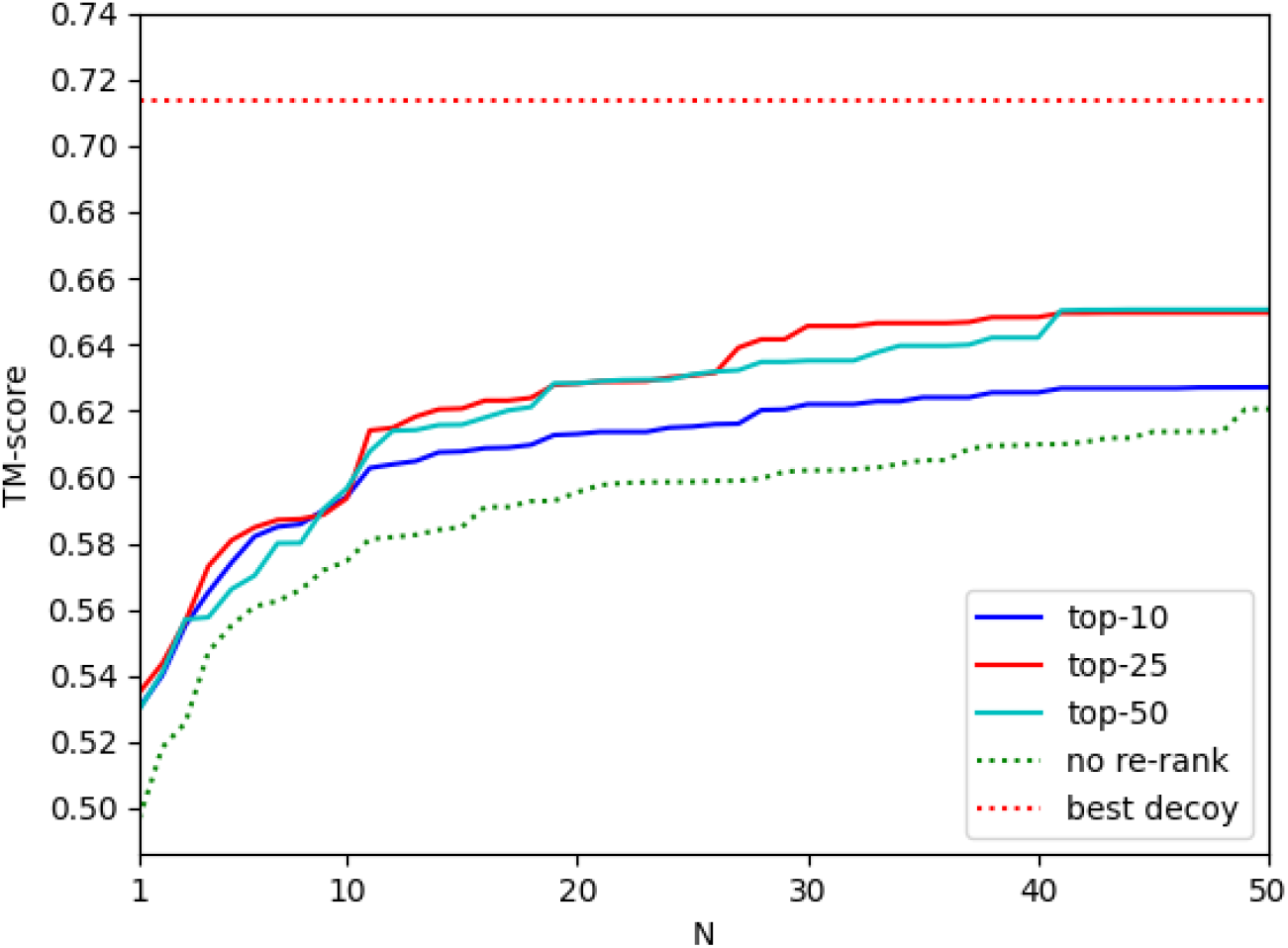
The average quality (measured by TMscore) of the selected decoys by top predicted contacts. The x-axis is the number of top decoys selected. In the legend, “top-10”, “top-25” and “top-50” represent that top 10, 25 and 50 predicted contacts are used to select docking decoys, respectively. “best decoy” indicates the quality of the best decoys generated by HDOCK.

## 4 Conclusion

We have presented an interfacial contact prediction method, GLINTER, that predicts inter-protein contacts by integrating attention information generated by protein language models and graph modeling of monomer (experimental and predicted) structures. The attention may capture some evolutionary and coevolutionary information encoded in MSA (multiple sequence alignment). We demonstrate that GLINTER outperforms existing methods and even if trained with experimental structures, it generalizes well to predicted structures. The interfacial contacts predicted by our method may help improve selection of docking decoys. Our ablation study shows that the attention information and structural features are complementary and important for interfacial contact prediction. The features used by GLINTER can be calculated very efficiently and GLINTER is applicable to both heterodimers and homodimers. Therefore, potentially GLINTER is applicable to the proteome-scale study of protein-protein interactions and complexes.

