## Supplemental File for "Deep graph learning of inter-protein contacts"

Ziwei Xie and Jinbo Xu*
Toyota Technological Institute at Chicago, Chicago, IL, USA

#### 1 Build a surface graph

To build a surface graph, we first use Reduce to add hydrogen atoms and then construct the triangulated surface of a protein structure using MSMS. MSMS may generate a large number of vertices on the triangulated surface. We use the “remove_closest” algorithm in the trimesh library[1] to sample a subset of vertices such that any two of them are at least 0.8Å away from each other. The cutoff is set to 0.8Å, because the number of remaining vertices does not change much when it is less than 0.8Å.

#### 2 Select a diverse set of sequences

To select a relatively small set of diverse sequences from an MSA, we use HHfilter[2] with -diff 200 and -cov 20 to filter MSAs[3]. Then we use Henikoff sequence weighting to rank the remaining sequences, and select at most 128 sequences for each concatenated MSA as the input of the MSA Transformer.

#### 3 MSA depth (Meff)

Given an MSA, we use CD-HIT[4,5] to cluster all the sequences in this MSA using 65% sequence identity as cutoff. The effective MSA depth is roughly defined as the number of clusters.

#### 4 Case study

We study an interesting target, T0997, where the ESM-Attention model and the Residue+Atom+Surface model perform much better than the Residue+Atom+Surface+ESM model. In Fig. S5, the cluster of contacts correctly predicted by the ESM-Attention model is different from the cluster correctly predicted by the Residue+Atom+Surface model, so we can hypothesize that the ESM-Attention model (MSA-based) and the Residue+Atom+Surface model (structure-based) focus on very different patterns in T0997. Therefore, it is possible that there are some structure patterns around the cluster correctly predicted by the ESM-Attention model that make the Residue+Atom+Surface+ESM model predictions misaligned with the ground truth contacts; see Fig. S4.

### 5 Tables

**Table S1.** The number of correct top-10,25,50 predictions by 3 different deep models on 32 dimer targets. “EA” denotes the ESM-Attention model. “RASE-N” denotes the Residue+Atom+Surface+ESM model using native structures as the inputs. “RAS-N” denotes the Residue+Atom+Surface model using native structures as the inputs.

| | 10 | | | 25 | | | 50 | | |
|---|---|---|---|---|---|---|---|---|---|
| Target | EA | RASE-N | RAS-N | EA | RASE-N | RAS-N | EA | RASE-N | RAS-N |
| H0957 | 0 | 6 | 0 | 1 | 10 | 3 | 1 | 16 | 7 |
| H0974 | 7 | 6 | 5 | 12 | 11 | 16 | 20 | 19 | 27 |
| H0986 | 3 | 0 | 0 | 10 | 0 | 0 | 14 | 0 | 0 |
| H1015 | 0 | 0 | 1 | 1 | 0 | 2 | 2 | 2 | 6 |
| H1017 | 0 | 8 | 3 | 0 | 17 | 7 | 0 | 24 | 12 |
| H1019 | 6 | 8 | 6 | 10 | 15 | 12 | 15 | 20 | 20 |
| H1045 | 7 | 7 | 0 | 12 | 14 | 0 | 22 | 17 | 0 |
| H1047 | 0 | 0 | 0 | 0 | 0 | 0 | 0 | 0 | 0 |
| H1065 | 2 | 5 | 3 | 4 | 12 | 5 | 10 | 20 | 12 |
| T0965 | 10 | 8 | 10 | 15 | 16 | 20 | 25 | 31 | 37 |
| T0966 | 0 | 0 | 6 | 0 | 0 | 8 | 0 | 0 | 12 |
| T0970 | 0 | 8 | 7 | 0 | 18 | 16 | 0 | 32 | 22 |
| T0973 | 0 | 9 | 7 | 1 | 20 | 14 | 8 | 37 | 23 |
| T0976 | 0 | 0 | 0 | 0 | 4 | 0 | 0 | 10 | 1 |
| T0983 | 0 | 2 | 1 | 0 | 4 | 1 | 1 | 7 | 6 |
| T0991 | 1 | 6 | 6 | 4 | 15 | 14 | 7 | 24 | 25 |
| T0997 | 5 | 0 | 2 | 8 | 1 | 7 | 12 | 4 | 19 |
| T0998 | 0 | 8 | 7 | 0 | 16 | 15 | 0 | 24 | 24 |
| T1000 | 6 | 3 | 0 | 11 | 8 | 0 | 23 | 14 | 5 |
| T1001 | 0 | 2 | 6 | 0 | 6 | 8 | 0 | 8 | 10 |
| T1003 | 10 | 10 | 7 | 24 | 24 | 12 | 47 | 41 | 23 |
| T1006 | 8 | 8 | 1 | 17 | 14 | 3 | 22 | 14 | 5 |
| T1010 | 0 | 9 | 6 | 0 | 14 | 8 | 0 | 23 | 11 |
| T1016 | 0 | 10 | 10 | 2 | 23 | 19 | 4 | 34 | 30 |

| | | | | | | | | | |
|---|---|---|---|---|---|---|---|---|---|
| T1018 | 2 | 0 | 0 | 8 | 0 | 0 | 14 | 0 | 0 |
| T1032 | 8 | 10 | 0 | 16 | 24 | 2 | 33 | 48 | 12 |
| T1038 | 0 | 2 | 5 | 0 | 2 | 7 | 0 | 7 | 13 |
| T1054 | 2 | 1 | 2 | 7 | 5 | 5 | 14 | 9 | 8 |
| T1078 | 4 | 6 | 10 | 9 | 14 | 19 | 12 | 21 | 27 |
| T1083 | 8 | 6 | 2 | 18 | 14 | 5 | 34 | 29 | 6 |
| T1084 | 8 | 10 | 5 | 19 | 25 | 9 | 33 | 50 | 13 |
| T1087 | 2 | 7 | 9 | 3 | 11 | 17 | 6 | 23 | 29 |

**Table S2**. Features used in spatial graphs, where L is the number of residues, N is the number of atoms, M is the number of sampled surface vertices, and E is the number of edges in an atom graph.

| Spatial Graph | Feature | Dimension |
|---|---|---|
| Residue graph | Positional Specific Scoring Matrix (PSSM) | L x 20 |
| | Residue solvent accessible surface areas (SASA) | L x 1 |
| | Amino acid types (AA) | L x 21 |
| | Normalized sequential positions | L x 1 |
| | CA coordinates | L x 3 |
| | CA-centered local reference frame (LRF) | L x 3 x 3 |
| Atom graph | Atom solvent accessible surface areas | N x 1 |
| | Atom chemical types | N x 10 |
| | Residue’s amino acid type | N x 21 |
| | Edge type | E x 1 |
| | Atom coordinates | N x 3 |
| Surface graph | Vertex coordinates | M x 3 |
| | Vertex normal vectors | M x 3 |

**Table S3.** Average interfacial contact precision of different deep learning models when experimental monomer structures are used. Column "D-cut" shows the distance cutoffs used to define graph edges. For example, "8,6,6" indicates that the residue graph, atom graph and surface graph use 8Å, 6Å, and 6Å to define edges, respectively.

| Model Settings | D-cut | 10 | 25 | 50 | L/10 | L/5 |
|---|---|---|---|---|---|---|
| ESM-Attention | | 30.93 | 26.50 | 23.69 | 29.38 | 28.02 |
| CNN+ESM-Attention | | 35.31 | 28.25 | 20.12 | 24.31 | 33.98 |
| Residue | 6 | 28.13 | 24.50 | 20.94 | 26.82 | 23.57 |
| | 8 | 30.63 | 26.25 | 22.94 | 30.06 | 27.01 |
| | 10 | 29.69 | 26.13 | 21.38 | 28.26 | 25.00 |
| Residue + ESM | 8 | 42.81 | 37.13 | 34.25 | 41.80 | 37.12 |
| Residue + Atom | 8,4 | 21.56 | 19.13 | 18.75 | 22.58 | 18.82 |
| | 8,6 | 32.50 | 28.50 | 23.06 | 31.85 | 27.39 |
| | 8,8 | 26.56 | 25.00 | 22.69 | 27.46 | 25.65 |
| | 8,10 | 29.69 | 25.50 | 22.19 | 26.55 | 26.75 |
| Residue + Atom + ESM | 8,6 | 39.06 | 34.00 | 31.88 | 36.26 | 33.84 |
| | 8,8 | 40.6 | 36.00 | 32.06 | 39.00 | 35.47 |
| Residue + Surface | 8,4 | 33.13 | 26.75 | 23.81 | 31.18 | 27.04 |
| | 8,6 | 33.75 | 28.25 | 25.63 | 33.69 | 27.58 |
| | 8,8 | 33.43 | 29.25 | 25.81 | 33.34 | 29.37 |
| | 8,10 | 32.50 | 29.25 | 26.19 | 32.36 | 27.90 |
| Residue + Surface + ESM | 8,6 | 44.06 | 37.88 | 33.88 | 41.38 | 37.92 |
| | 8,8 | 39.69 | 34.50 | 31.31 | 36.95 | 36.11 |
| Residue + Atom + Surface | 8,6,6 | 39.69 | 31.75 | 27.81 | 35.97 | 31.53 |
| Residue + Atom + Surface + ESM | 8,6,6 | **51.56** | **44.63** | **38.00** | **50.36** | **44.43** |

**Table S4.** Average interfacial contact precision of different deep learning models when monomer structures are predicted by AlphaFold. Column "D-cut" shows the distance cutoffs used to define graph edges.The performance of the ESM-Attention model in Tables S3 and S4 is not the same because the predicted and experimental monomer structures do not have exactly the same set of residues.

| Model Settings | D-cut | 10 | 25 | 50 | L/10 | L/5 |
|---|---|---|---|---|---|---|
| ESM-Attention | | 29.06 | 25.63 | 22.94 | 27.26 | 26.97 |
| CNN+ESM-Attention | | 21.56 | 19.50 | 17.75 | 19.59 | 18.67 |
| Residue | 6 | 21.56 | 18.75 | 16.94 | 22.03 | 18.68 |
| | 8 | 26.56 | 23.62 | 19.50 | 25.93 | 22.07 |
| | 10 | 22.50 | 20.13 | 18.00 | 23.17 | 20.75 |
| Residue + ESM | 8 | 33.44 | 30.00 | 29.06 | 33.21 | 29.69 |
| Residue + Atom | 8,4 | 23.44 | 21.50 | 18.06 | 24.45 | 19.66 |
| | 8,6 | 26.25 | 21.25 | 17.69 | 23.91 | 20.07 |
| | 8,8 | 26.25 | 23.63 | 18.56 | 28.72 | 22.70 |
| | 8,10 | 25.31 | 23.13 | 19.44 | 25.97 | 23.17 |
| Residue + Atom + ESM | 8,6 | 35.31 | 30.25 | 27.94 | 31.86 | 28.98 |
| | 8,8 | 30.62 | 28.25 | 26.63 | 30.21 | 27.13 |
| Residue + Surface | 8,4 | 24.06 | 19.75 | 17.19 | 23.22 | 20.54 |
| | 8,6 | 22.19 | 19.75 | 18.81 | 21.27 | 17.45 |
| | 8,8 | 27.81 | 23.88 | 20.18 | 25.19 | 22.45 |
| | 8,10 | 27.50 | 24.88 | 21.19 | 26.52 | 24.48 |
| Residue + Surface + ESM | 8,6 | 36.56 | 32.75 | 29.75 | 34.51 | 31.81 |
| | 8,8 | 34.69 | 30.5 | 28.25 | 32.87 | 30.24 |
| Residue + Atom + Surface | 8,6,6 | 29.38 | 23.88 | 20.25 | 27.07 | 24.06 |
| Residue + Atom + Surface + ESM | 8,6,6 | **37.81** | **34.38** | **31.31** | **37.23** | **32.94** |

### 6 Figures

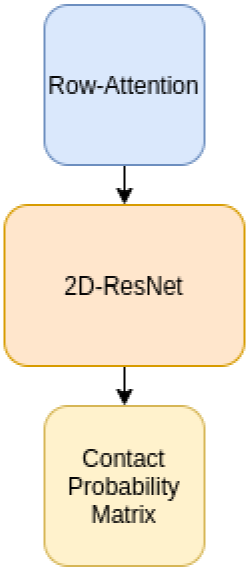


**Fig. S1.** The ESM-Attention model

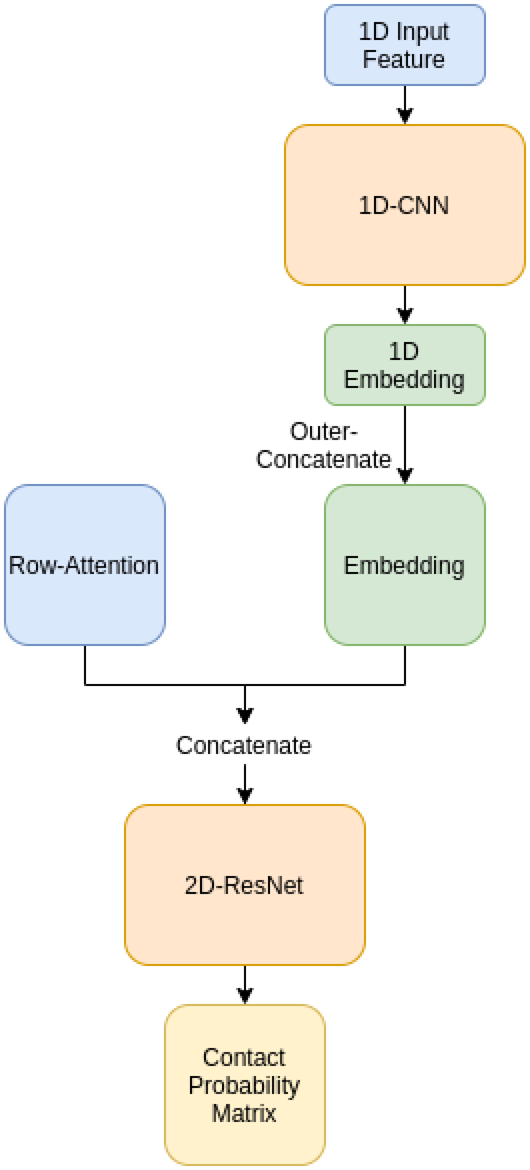


**Fig. S2.** CNN+ESM-Attention model

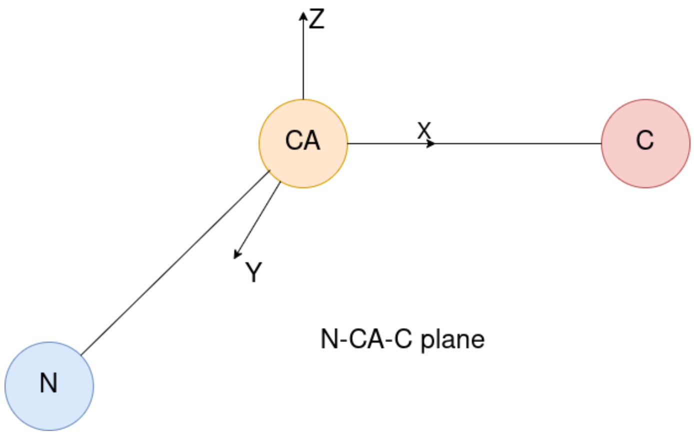


**Fig. S3.** Local reference frame. The CA-C bond is used as the x-axis, the z-axis is perpendicular to the N-CA-C plane, and the y-axis is the cross-product of x-axis and z-axis.

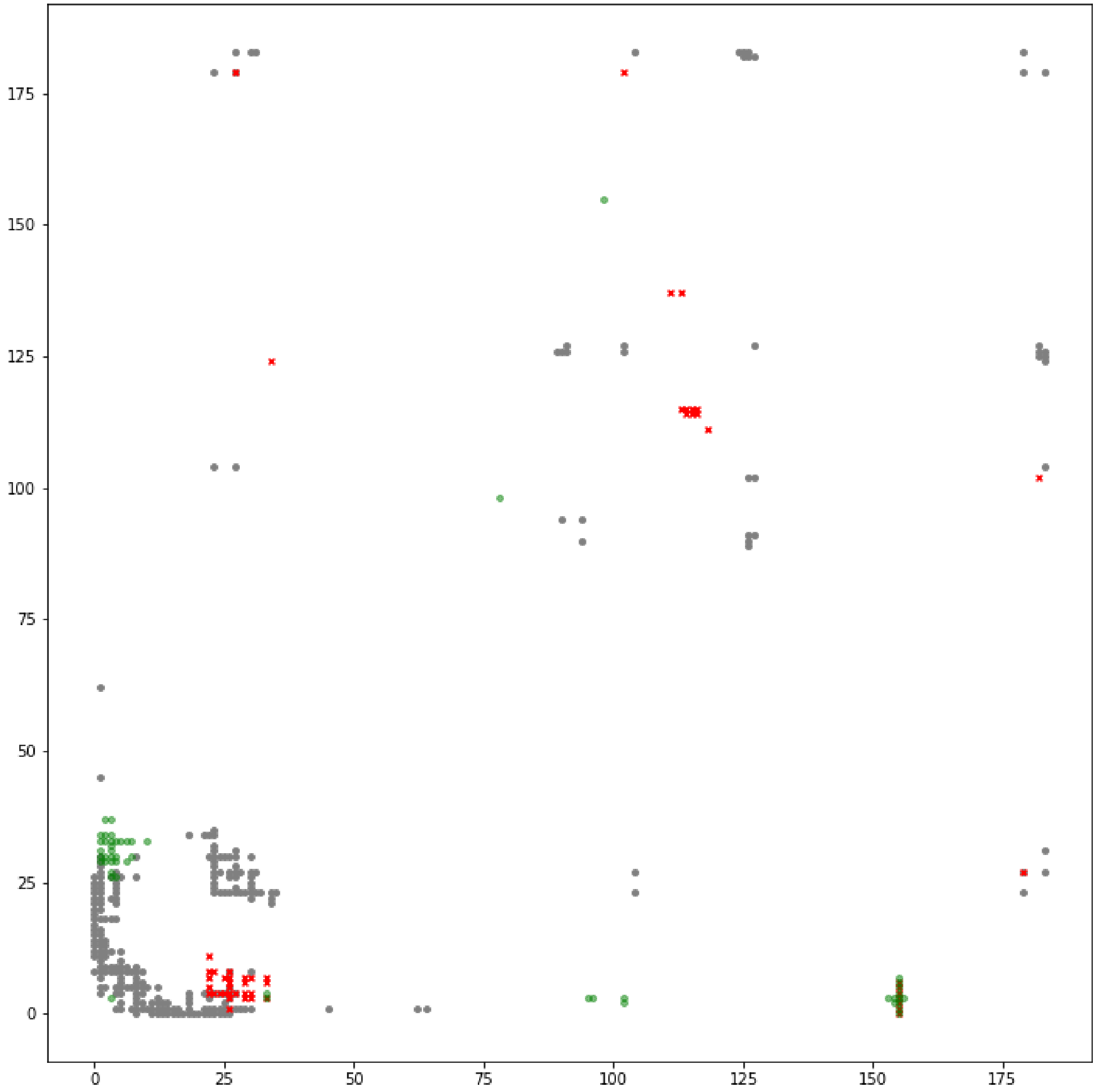


**Fig. S4.** Interfacial contact prediction of T0997. The grey dots indicate the ground truth contact, the red crosses indicate the top 50 predictions of the ESM-Attention model, and the green dots indicate the top 50 predictions of the Residue+Atom+Surface+ESM (native) model.

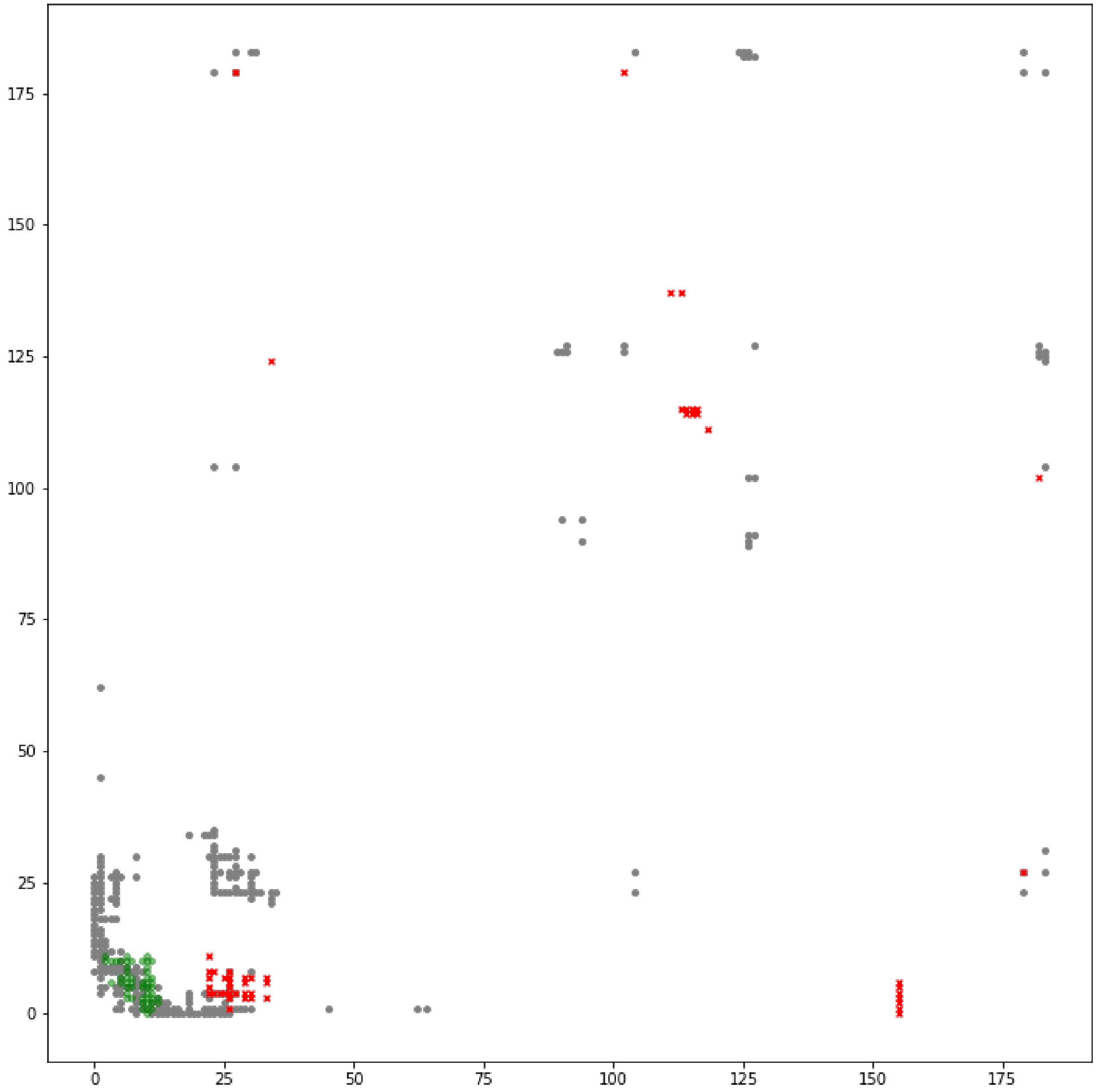


**Fig. S5.** Interfacial contact prediction of T0997. The grey dots indicate the ground truth contacts, the red crosses indicate the top 50 predictions of the ESM-Attention model, and the green dots indicate the top-50 predictions of the Residue+Atom+Surface (native) model.

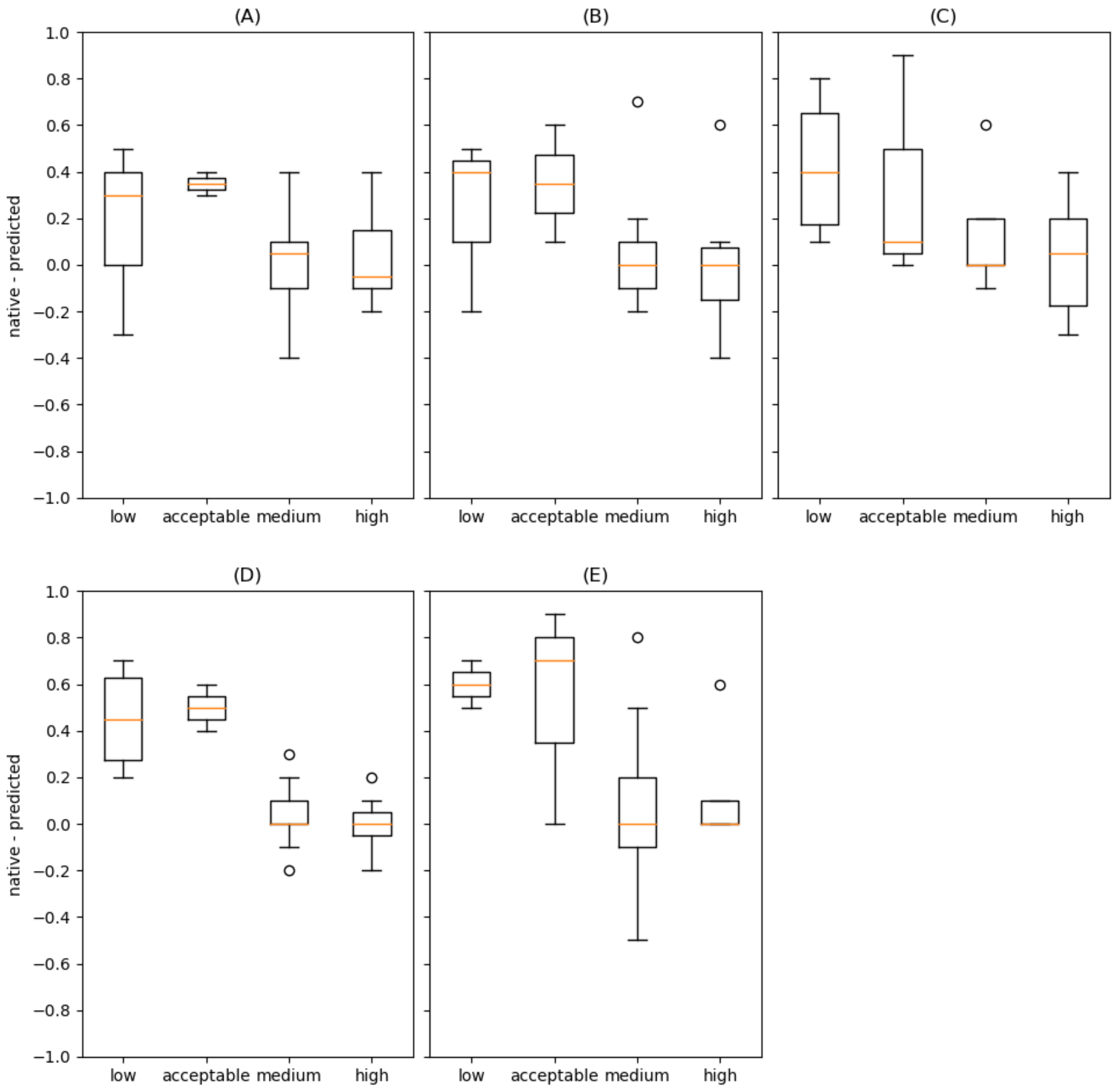


**Fig. S6.** The x-axis is the TMscore of the predicted monomer structures. The y-axis is the difference of the top 10 precision resulting from the experimental and predicted monomer structures. (A) “Residue, D-cut=8”. (B) “Residue + Atom, D-cut=8,6” (C) “Residue + Surface, D-cut=8,6” (D) “Residue + Atom + Surface, D-cut=8,6,6” (E) “Residue + Atom + Surface + ESM, D-cut=8,6,6”. In all the box plots, the upper edge of the box is the third quartile (Q3), and the lower edge of the box is the first quartile (Q1), the orange line is the median, the upper cap is the highest datum below Q3 + 1.5(Q3 - Q1), and the lower cap is the lowest datum above Q1 - 1.5(Q3-Q1).

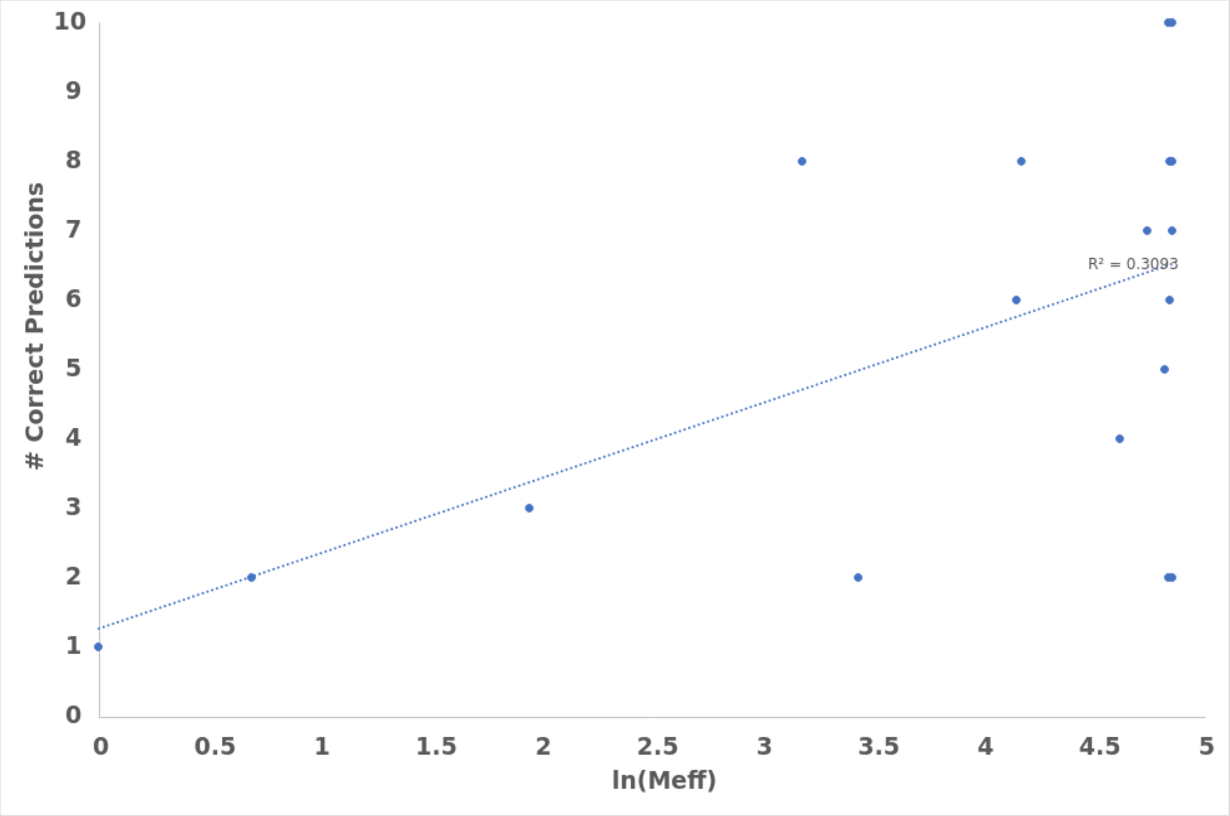


**Fig. S7.** Correlation between ln(Meff) (x-axis) and the number of correct top-10 predictions (y-axis) of the ESM-Attention model. The targets without correct top-10 predictions are excluded. ($R^2 = 0.3093$)
